# Exploring repositioning opportunities and side-effects of statins: a Mendelian randomization study of HMG-CoA reductase inhibition with 55 complex traits

**DOI:** 10.1101/170241

**Authors:** SO Hon-Cheong, Carlos Kwan-long Chau, Kai Zhao

**Author notes:** Correspondence to: Hon-Cheong SO.

## Abstract

Statin is one of the most commonly prescribed medications worldwide. Besides reduction of cardiovascular risks, statins have been proposed for the prevention or treatment of other disorders, but results from clinical studies are mixed. There are also controversies concerning the adverse effects caused by statins.

In this study we employed a Mendelian randomization (MR) approach across a wide range of complex traits to explore repositioning opportunities and side-effects of statins. MR is analogous to a “naturalistic” randomized controlled trial (RCT), which is much less susceptible to confounding and reverse causation as compared to observational studies.

We employed two genetic instruments (rs12916 and rs17238484) in the *HMGCR* gene which have been shown to provide reliable estimates of the risk of statins on type 2 diabetes and weight gain. We observed in the single- and joint-SNP analysis that low density lipoprotein cholesterol (LDL-C) reduction from HMG-CoA reductase inhibition results in increased depressive symptoms. This finding appeared to be supported by nominally significant results of raised major depression risk in single-SNP MR analysis of rs17238484, and analyses using LDL-C as the exposure. Several other outcomes also reached nominal significance (*p* < 0.05) in single- or joint-SNP analyses; for example, we observed causal associations of LDL-C lowering from HMG-CoA reductase inhibition with *reduced* risks of schizophrenia, anorexia nervosa, Alzheimer disease, Parkinson disease, as well as increased forearm bone mineral density, sleep duration and extreme longevity (highest *q-value* = 0.289). We also found evidence of casual relationships of LDL-C levels with schizophrenia, anorexia, sleep duration and longevity, following the same association directions as in analyses of *HMGCR* variants. These findings were at least partially supported by previous clinical studies. We did not observe associations with cognitive test profiles, renal outcomes, autoimmune diseases or cancers. While MR has its limitations and our findings remain to be confirmed in further studies, this work demonstrates the potential of a phenome-wide approach to reveal novel therapeutic indications and unknown drug side-effects.

## INTRODUCTION

With the rising cardiovascular disease burden over the world, statins have become one of the most commonly prescribed classes of medications. For example, it was estimated that in the US over 25% (30 millions) of people were taking statins from 2005 to 2008^1^. Statins act on the 3-hydroxy-3-methyl-glutaryl-coenzyme A (HMG-CoA) reductase to reduce cholesterol synthesis, and there exist strong evidence that statins reduce cardiovascular risks^2-4^. Statins however are associated with side-effects such as myopathy, raised liver enzymes and increased risk of type 2 diabetes and probably hemorrhagic strokes^5^. There are also reports of statins being associated with other adverse effects, such as muscle-related symptoms other than myopathy^6^, cognitive decline^7-9^, mood changes or aggression^10,11^ and cataracts^12^, but there is yet no conclusive evidence. On the other hand, statins have also been proposed as a treatment for other non-cardiovascular traits, for example in cancers^13^, neurological diseases^14^, infectious diseases^15^ and psychiatric disorders^14^. Since statins are very widely prescribed, understanding its side-effects as well as non-cardiovascular benefits and the corresponding mechanisms is of great public health importance. Also given the escalating cost in new drug development, repositioning of a relatively cheap medication like statin represents a cost-effective way of finding novel therapeutics.

Recently a few studies have been using a new approach known as Mendelian randomization (MR)^16^ to explore the efficacy or side-effects of medications ^1,17-21^. The principle is that genetic variants in the gene encoding the drug target act as “instruments” to reflect the actual drug action. A person having the lipid-lowering allele (or allelic score) at the relevant locus is analogous to receiving a lipid-lowering drug, and vice versa. The random allocation of alleles at conception is analogous to randomization in clinical trials. MR, when compared to conventional clinical studies, are much less susceptible to confounding bias and problems of reversed causality^22^. By studying genetic variations at the drug target, we may also infer whether the effects or side-effects are at least partially due to “on-target” mechanisms. MR therefore represents a promising approach to explore new indications and unintended adverse effects of drugs, as reported in a few recent studies. For example, Swerdlow et al. examined common variants in the *HMGCR* gene and found that the low density lipoprotein cholesterol (LDL-C) lowering allele is associated with higher body weight, waist circumference, insulin and glucose levels, as well as heightened type 2 diabetes risk. Remarkably, the results from this MR analysis are highly concordant with a meta-analysis of randomized controlled trial (RCT) of 129710 individuals^1^. This study also demonstrated that raised diabetic risk is at last partially attributed to “on-target” effects of statins^1^. Adopting a similar approach, Lotta et al. showed that SNPs in the *NPC1L1* and *HMGCR* genes (proxy for actions of ezetimibe and statins respectively) are both associated with elevated diabetic risks^21^. Another two MR studies on PCSK9 inhibition demonstrated similar findings of raised diabetic risks^18,19^.

The MR approach can be extended to screen for associations of a larger variety of outcomes, also referred to as a “phenome-wide” scan. For instance, Millard et al. reported a phenome-wide MR study on BMI with 172 phenotypic outcomes, and found associations with cardiometabolic traits as well as novel associations with global self-worth score^23^. Another recent study investigated the effect of a loss-of-function variant in *PLA2G7* encoding lipoprotein-associated phospholipase A2 (Lp-PLA2), a drug target for atherosclerosis-related diseases, across 41 different non-vascular outcomes in the China Kardoorie Biobank^24^. Overall no significant associations were found, implying a lack of major side-effects but also limited potential for repurposing^24^. A recent commentary nicely summarizes the potential of using MR and a “phenome-wide” approach to facilitate drug discovery and reveal unknown drug side-effects^25^.

In this study, we employ the principle of Mendelian randomization to explore repositioning opportunities and adverse effects of statins. Using variants of the *HMGCR* gene as instruments, we study the associations with up to 55 somatic and psychiatric traits, mainly on non-cardiovascular outcomes.

## METHODS

### Genetic instruments for effects of statins

We followed a previous landmark MR study which investigated the effects of HMG-CoA reductase inhibition on body-weight and type 2 diabetes risks^1^. We used two SNPs (rs17238484 and rs12916) in the *HMGCR* gene as instruments that have been shown to provide reliable estimates of the risk of statins on type 2 diabetes and weight gain. The variant rs17238484 was used for the main analysis in Swerdlow et al., and the LDL-lowering G allele was associated with higher waist circumference, weight, insulin and glucose concentrations and type 2 diabetes risk^1^. The effects were consistent with (and of comparable magnitude to) meta-analysis of RCTs covering 129170 participants. The other SNP allele rs12916-T had similar effects in general. As reported by Swerdlow et al., rs12916-T is also associated with significantly lower expression of *HMGCR* in the liver (*p* = 1.3e-5) but has no associations with expressions of neighboring genes^1^. It was shown that this SNP likely drives the shared association between expression QTL and LDL-C levels^1^. In addition, the two SNPs achieved genome-wide significance (*p* < 5e-8) in the Global Lipid Genetics Consortium (GLGC) meta-analysis^26^ (Table 1). The effect sizes and standard errors of rs17238484 and rs12916 were extracted from meta-analysis results of the GLGC Metabochip studies. Note that the original GLGC study use inverse normal transformed trait values as the outcome variable, hence the coefficient estimate (approximately) corresponds to one SD change (~ 38.7 mg/dl or ~ 1 mmol/L) of LDL-C level per unit change in allelic count.

**Table 1.** SNPs used as genetic instruments in Mendelian randomization analysis

| chr | SNP | effect_allele<br>(lipid-lowering) | other_allele | beta | se | *p* |
| --- | --- | --- | --- | --- | --- | --- |
| 5 | rs12916 | T | C | 0.0695 | 0.0053 | 1.79E-34 |
| 5 | rs17238484 | G | T | 0.0627 | 0.0062 | 1.47E-21 |

Effect sizes are extracted from meta-analysis performed by the Global Lipid Genetics Consortium. Beta: regression coefficient; se, standard error; eaf, effect allele frequency; beta\_joint and se\_joint, coefficient and standard error estimate from COJO-joint analysis.

As the functional significance of *HMGCR* variants has not been fully elucidated (we do not know exactly which SNP or SNP combinations may represent the best proxy for the action of statins), we performed both single-SNP analyses (as in Swerdlow et al.^1^) and a combined analysis of both SNPs. Note that as both SNPs are located in the *HMGCR* gene (cis-acting variants), they are likely to exert their effects via pathways related *HMGCR,* instead of through other mechanisms. Therefore we believe that horizontal pleiotropy is unlikely (https://www.ncbi.nlm.nih.gov/pmc/articles/PMC5100611/), although the small number of genetic markers does not allow more quantitative assessment of pleiotropy by methods like Egger regression or the weighted median approach.

### Outcome data

We made use of the MR-base database and R package TwoSampleMR which collects GWAS summary statistics from mostly publicly available sources^27^. A small percentage of the data available at MR-base were not openly available but obtained through communication with individual investigators. We performed analyses on several categories of complex traits or diseases as follows:

1. Psychiatric disorders or traits including anorexia nervosa, autism, bipolar disorder, bulimia nervosa, depressive symptoms, major depressive disorder (MDD), schizophrenia and sleep duration (for;
2. Neurological disorders including Alzheimer’s disease, amyotrophic lateral sclerosis and Parkinson’s disease;
3. Renal traits or disorders including chronic kidney disease, microalbuminuria, IgA nephropathy; eGFR (creatinine) and urinary albumin to creatinine ratio (UACR) within diabetic subjects and within non-diabetic subjects;
4. Cancers including melanoma, pancreatic cancer, neuroblastoma, lung adenocarcinoma, squamous cell lung cell and lung cancer (irrespective of subtypes);
5. Autoimmune or inflammatory conditions including celiac disease, asthma, eczema, gout, Crohn’s disease, ulcerative colitis, inflammatory bowel disease (combined), multiple sclerosis, rheumatoid arthritis, systemic lupus erythematosus;
6. Bone mineral density (BMD) measures including femoral neck, forearm and lumbar spine BMDs;
7. Hematological traits including haemoglobin (Hb) concentration, mean cell Hb, mean cell Hb concentration, mean cell volume, packed cell volume, red blood cell count, platelet count and mean platelet volume;
8. Neurocognitive test profiles including 2-choice, 4-choice and 8-choice reaction times, digit symbol, G speed factor, inspection time, simple reaction time and symbol search. This category is included due to previous reports of possible links between statin use and cognitive deficits (e.g. ref. ^8,9^);
9. Aging and longevity related traits including age of parents’ death and top 1% survival in parents, which serve a proxy measure of longevity in the individual. The original study by Pilling et al.^28^ explains the advantages of such an approach instead of recruiting very old cases and younger controls. An exact causal inference was not attempted here but the genetic variants or genetic score of the offspring can be regarded to reflect the genetically determined LDL-C-level (due to HMG-CoA reductase inhibition) of the parents. We aimed to test for a genetic (or polygenic) association with longevity similar to Marioni et al.^29^. The effect size estimate is likely to be biased downwards as we are over-estimating the correlation between the genetic instruments and the risk factor (the denominator of an MR estimate).

A full list of references is included in supplementary information.

### Statistical analysis

Mendelian randomization was performed with the Wald ratio test for individual SNP analyses and the inverse variance weighted (IVW) approach for analyses involving more than one SNP, which are default methods employed in TwoSampleMR^27^. If a SNP was not available in the outcome GWAS, we allowed using a “proxy SNP” provided the r-squared was at least 0.8 with the requested SNP. LD information was taken from the 1000 Genome European samples.

For the joint analyses of rs12916 and rs17238484, we accounted for the correlation using the method described in Burgess et al.^30^. Briefly, assume 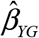 to be the vector of estimated regression coefficients when the outcome is regressed on genetic instruments and *σ_YG_* to be the corresponding standard errors (SE), and 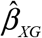 to be the estimated coefficients when the risk factor is regressed on the genetic instruments with SE *σ_XG_*. We also assume the correlation between two genetic variants G_1_ and G_2_ to be *ρ_G1G2_*, and ∑_*G*_1_*G*_2__ = *ρ*_*G*_1_*G*_2__ *σ*_*YG*_1__ *σ*_*YG*_2__.

The estimate from a weighted generalized linear regression can be formulated by

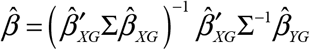

with SE

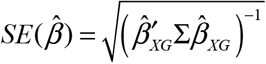

We also performed the joint analysis by maximum likelihood estimation ^31^and the results were largely similar.

For a significant association of statins with a measured outcome, the association can be due to a general effect on LDL-C lowering, or more specifically due to HMG-CoA reductase inhibition. We therefore also examined the causal effects of LDL-C levels on outcomes having at least one nominally significant result. Totally 11 outcomes were studied. Genetic instruments for LDL-C were chosen according to the GLGC GWAS meta-analysis results^26^. We included two sets of GWAS summary statistics, one from Metabochip and the other from joint GWAS and Metabochip analyses. (Note that in the study on *HMGCR* variants, we only considered results from Metabochip as one of the variants (rs17238484) was not reported in the latter joint analyses.)

We then performed further analyses to examine the causal effects of LDL-C on various outcomes. We again adopted the methodology by Burgess et al.^30^ which is able to account for correlations among the instrument genetic variants. The method was based on formula (1) as described above. As remarked by the authors, inclusion of a bigger panel of variants in partial LD may enable higher variance to be explained, thus improving the power of MR. Including “redundant” SNPs in addition to the causal variant(s) do not improve power but also will not invalidate the results. However, including too many variants with high correlations may also render the MR estimates unstable^32^. But there is a lack of strong theoretical justifications on the choice of a single specific r^2^ threshold for analysis. We therefore attempted a range of thresholds, but controlled for multiple testing with FDR. We conducted MR analysis at four levels of r^2^ threshold (0.05, 0.1, 0.15, 0.2) within a 10Mb region, using the IVW and Egger approach implemented in the R package “MendelianRandomization”, and accounted for SNP correlations at the same time. The intercept from Egger regression was used to assess whether there is significant directional pleiotropy (if *p* < 0.05). We primarily considered the results from IVW approach if there is no significant directional pleiotropy, otherwise the estimates from Egger regression were presented.

Multiple testing of different thresholds and the two GWAS datasets was corrected by the false discovery rate (FDR) approach with the Benjamini-Hochberg method^33^, which controls the expected proportion of false discoveries. A *q*-value represents the minimum FDR at which an observed statistic can be called significant. In this study we set a primary FDR threshold of 0.05 to declare significance; results with 0.05 < *q* < 0.2 were considered suggestive associations. Some of the test results were positively correlated due to overlapping genetic variants, but the FDR approach is also valid under positive (regression) dependencies^34^. Correction by Bonferroni approach will be overly conservative due to correlations in hypotheses, in addition the Bonferroni approach controls the probability of making any single false discovery (to be < 0.05 usually), which have been argued to be too conservative in many settings^35^. Here our aim is to discover potential casual relationships of statins or LDL-C with health traits, which are to be followed up or validated in further studies; controlling the proportion of false discoveries would fit this purpose. Moreover, the interpretation of Bonferroni-corrected tests depends on the number of tests performed, which can be subject to manipulations such as selective reporting to achieve the desired significance threshold^35^.

## RESULTS

The effect size estimates of the SNPs rs17238484 and rs12916 from GLGC and COJO-joint analysis are listed in Table 1. The alleles associated with *lower* LDL-C levels are designated as the effect alleles. Note that the coefficient estimates refer to the effect of *LDL-C lowering* due to HMG-CoA reductase inhibition.

### Single-SNP analyses

For single-SNP analysis of rs17238484 (Table 2), a number of outcomes showed nominal significance, including anorexia nervosa (beta = -1.46, OR = 0.231, 95% CI = 0.068 to 0.786, p = 0.019), major depressive disorder (beta = 0.863, OR = 2.37, 95% CI = 1.06 to 5.34, p = 0.037), schizophrenia (beta = -0.496, OR = 0.609, 95% CI = 0.413 to 0.897, p = 0.012), sleep duration (beta = 0.186, 95% CI = 0.0378 to 0.334, p = 0.014), Alzheimer disease (beta = -0.797, OR = 0.450, 95% CI = 0.253 to 0.801, p = 6.58e-3), Parkinson disease (beta = -1.779, OR = 0.169, 95% CI = 0.035 to 0.803, p = 0.025), celiac disease (beta = 1.13, OR = 3.10, 95% CI = 1.04 to 9.18, p = 0.042) and forearm bone mineral density (beta = 0.719, 95% CI = 0.125 to 1.31, p = 0.018). The corresponding q-values ranged from 0.209 to 0.289. The results suggest possible beneficial effects of genetically lower LDL-C from HMG-CoA reductase inhibition on the aforementioned diseases, except for MDD and celiac disease where the effects were towards increased risks. As an example, the results suggest that one SD (~ 38.7 mg/dl or ~ 1 mmol/L) decrease in LDL-C genetically due to HMG-CoA reductase inhibition leads to an increased risk of MDD by ~ 2-fold (OR = 2.37) and higher forearm bone density by 0.719 SD units. The other results may be interpreted in a similar manner.

**Table 2.**
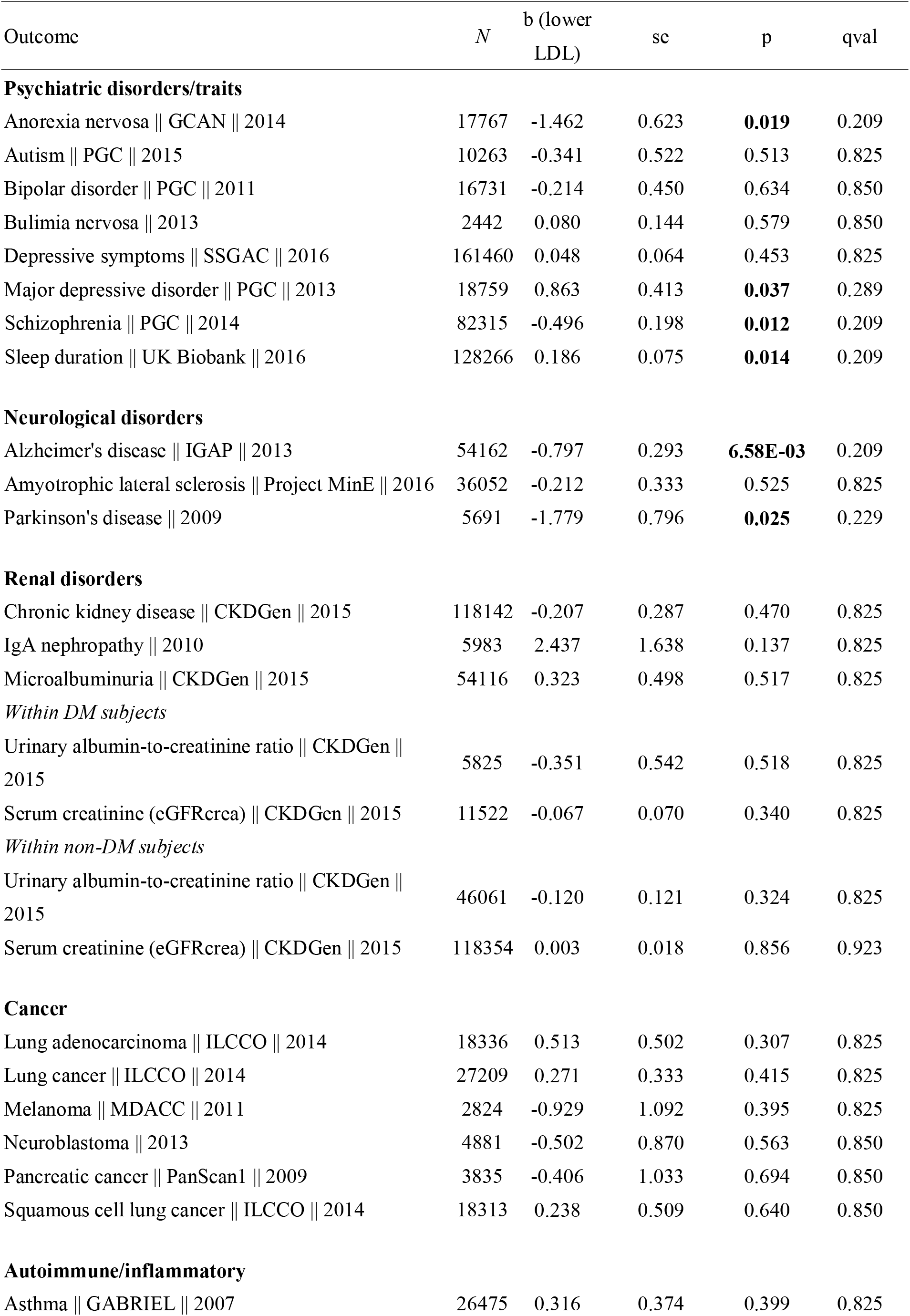
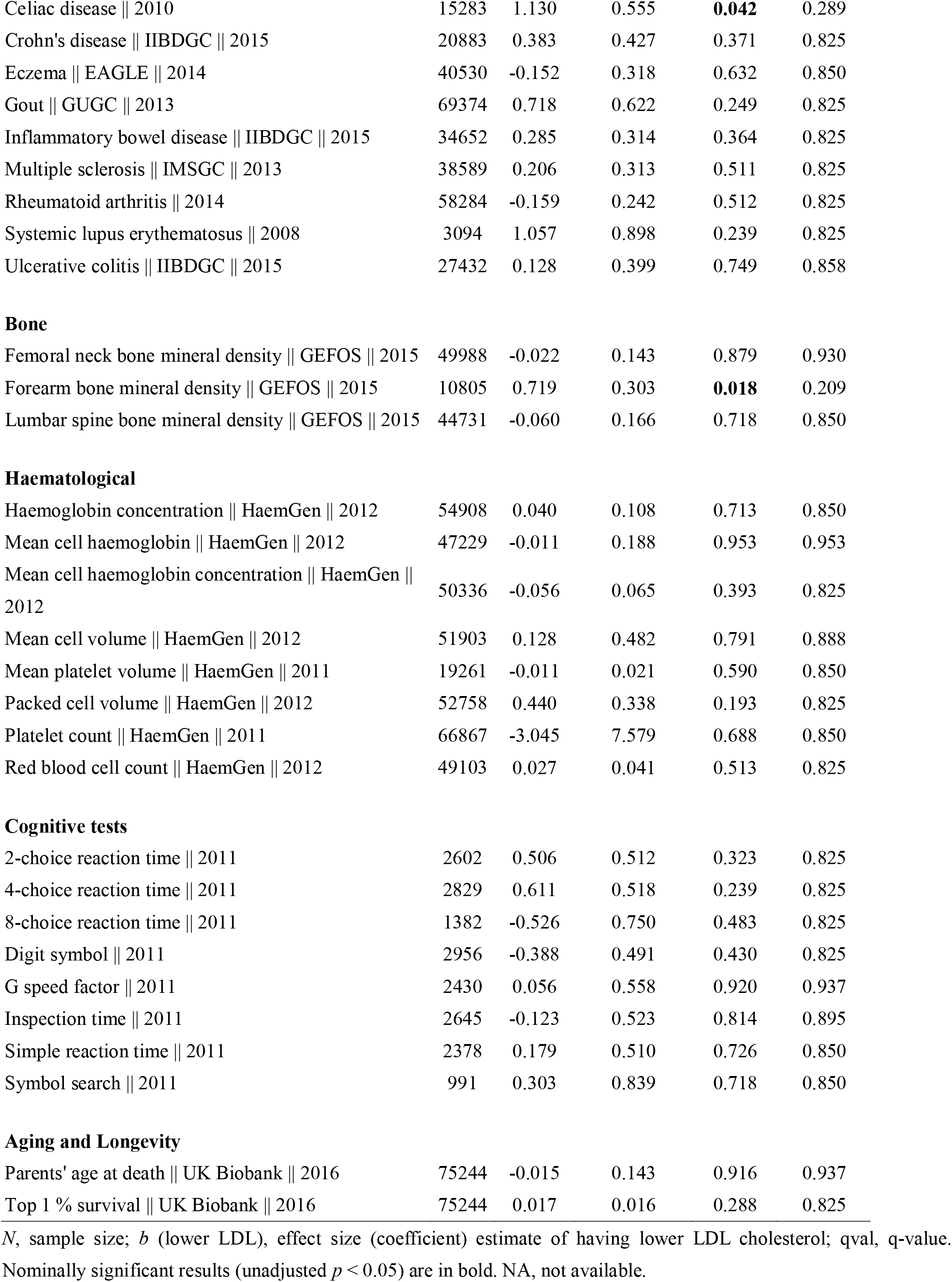
Mendelian randomization analysis of *HMGCR* variant rs1723 8484

Single-SNP analysis of rs12916 yielded three results with unadjusted *p* < 0.05 (Table 3), including depressive symptoms (beta = 0.129, 95% CI = 0.045 to 0.214, p = 2.70e-3), platelet count (beta = -12.6, 95% CI = 0.458 to 24.7, p = 0.042) and top 1% survival (parental survival as proxy) (beta = 0.025, OR = 1.025, 95% CI = 1.00 to 1.05, p = 0.048). Of note, there was a trend towards significance for anorexia nervosa (p = 0.063); this trait also showed significant casual associations in single-SNP analysis of rs17238484.

**Table 3.**
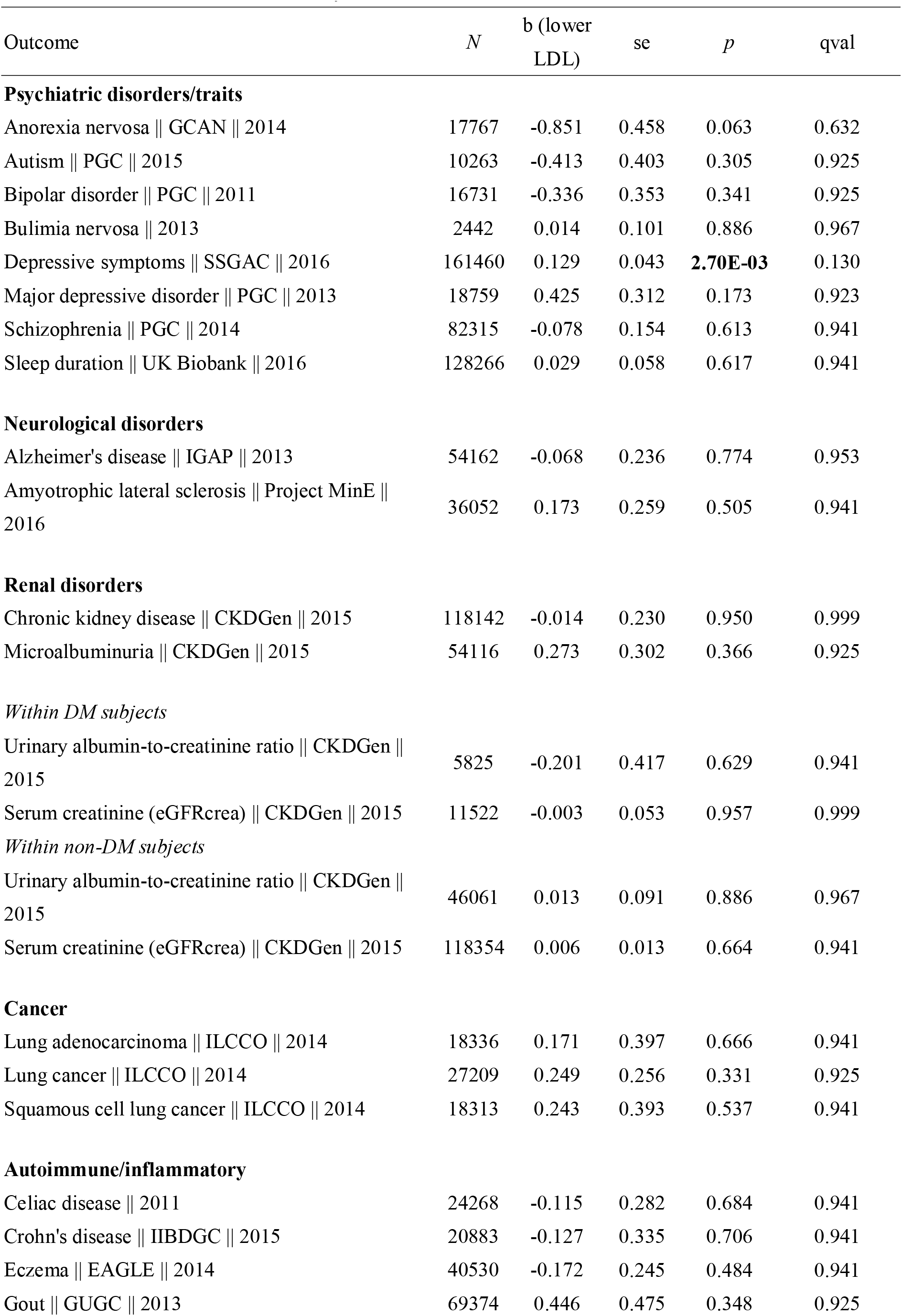
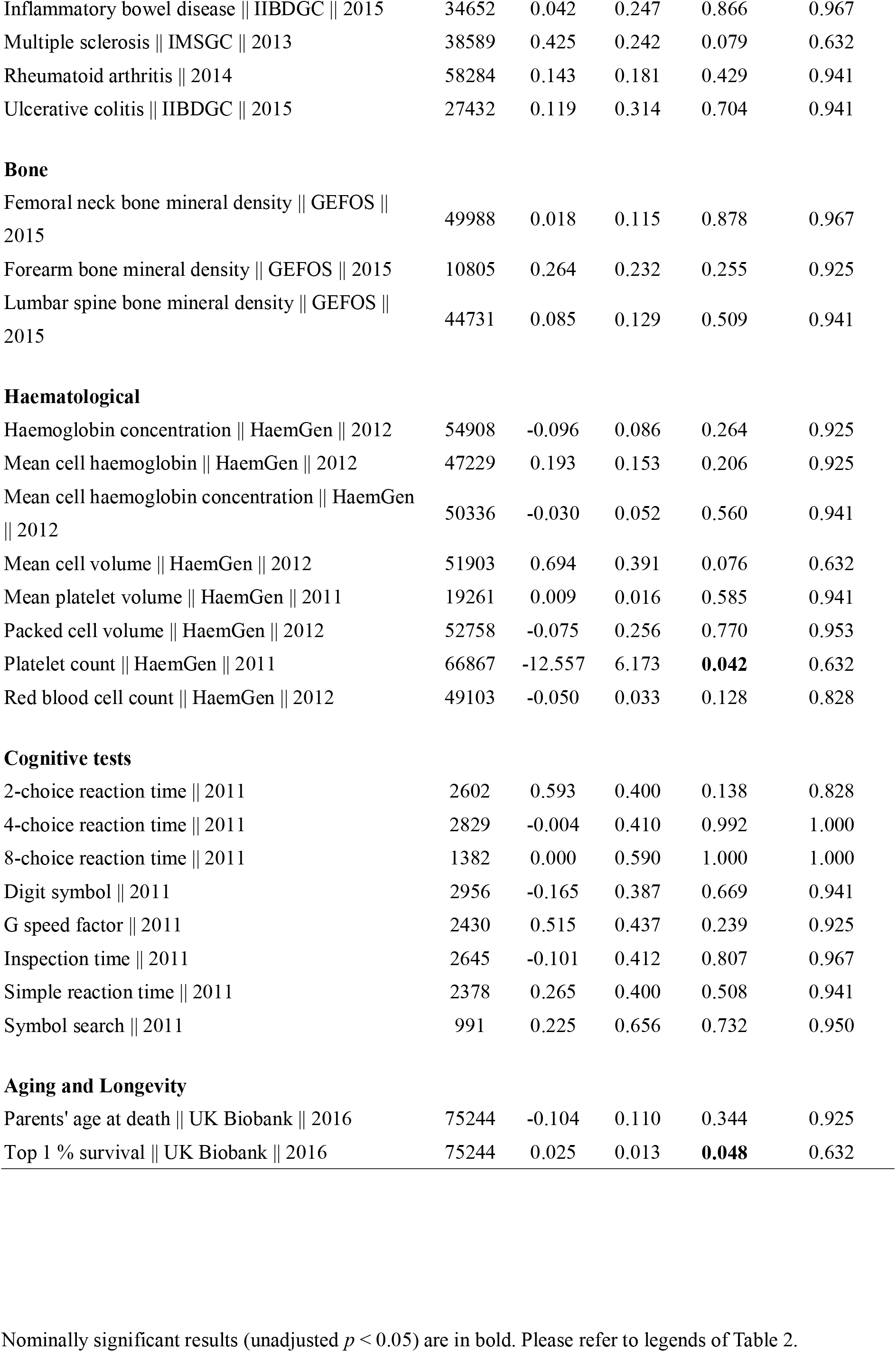
Mendelian randomization analysis of *HMGCR* variant rs12916

### Combined analysis of rs17238484 and rs12916

For the combined analysis of rs17238484 and rs12916, two outcomes achieved nominal significance (p < 0.05), including depressive symptoms (beta= 0.129, 95% CI = 0.045 to 0.214, *p* = 2.77E-3, *q* = 0.127) and anorexia nervosa (beta = -0.901, OR = 0.406, 95% CI = -1.797 to -0.006, p = 0.048, q= 0.822). Top 1% survival showed a trend towards significance (beta = 0.024, OR = 1.024, 95% CI = 0.999 to 1.049, p = 0.056, q = 0.822) (Supplementary Table 1).

### MR analysis of LDL-C with selected outcomes

We also performed MR analysis with LDL-C as the exposure for selected outcomes with nominal significant results (Table 4). Of note, the directions of associations from this analysis are fully concordant with the joint- and single-SNP analyses of *HMGCR,* except for celiac disease.

**Table 4.** Mendelian randomization analysis using LDL-C as exposure

| Outcome | Sample size | $r^2$ thres | Metabochip /joint | Pleiotropy pval | Beta (lower LDL) | SE | 95% CI Lower | 95% CI Upper | pval (Egger if pleiotropy +ve) | MR method | qval |
| --- | --- | --- | --- | --- | --- | --- | --- | --- | --- | --- | --- |
| Anorexia nervosa \|\| GCAN \|\| 2014 | 17767 | 0.15 | Metabochip | 0.260 | -0.202 | 0.078 | -0.354 | -0.049 | 9.61E-03 | IVW | *7.69E-02* |
| Depressive symptoms \|\| SSGAC \|\| 2016 | 161460 | 0.1 | jointGWAS | 0.161 | 0.028 | 0.006 | 0.015 | 0.040 | 1.04E-05 | IVW | **8.32E-05** |
| Major depressive disorder \|\| PGC \|\| 2013 | 18759 | 0.15 | jointGWAS | 0.671 | 0.140 | 0.036 | 0.070 | 0.210 | 9.31E-05 | IVW | **7.45E-04** |
| Schizophrenia \|\| PGC \|\| 2014 | 82315 | 0.2 | jointGWAS | 0.123 | -0.041 | 0.014 | -0.067 | -0.014 | 2.81E-03 | IVW | **1.96E-02** |
| Sleep duration \|\| UK Biobank \|\| 2016 | 128266 | 0.2 | Metabochip | 0.820 | 0.034 | 0.007 | 0.021 | 0.047 | 2.36E-07 | IVW | **1.89E-06** |
| Parkinson's disease \|\| NA \|\| 2009 | 5691 | 0.1 | Metabochip | 0.308 | 0.273 | 0.162 | -0.045 | 0.590 | 9.22E-02 | IVW | 5.16E-01 |
| Platelet count \|\| HaemGen \|\| 2011 | 66867 | 0.2 | jointGWAS | 0.930 | -3.274 | 0.846 | -4.931 | -1.616 | 1.08E-04 | IVW | **8.64E-04** |
| Forearm bone mineral density \|\| GEFOS \|\| 2015 | 10805 | 0.15 | jointGWAS | 0.927 | -0.048 | 0.024 | -0.096 | 0.000 | 5.00E-02 | IVW | 3.49E-01 |
| Celiac disease \|\| NA \|\| 2010 | 15283 | 0.15 | jointGWAS | 0.001 | -0.289 | 0.096 | -0.477 | -0.102 | 2.52E-03 | Egger | **2.02E-02** |
| Top 1 % survival \|\| UK Biobank \|\| 2016 | 75244 | 0.2 | jointGWAS | 8.78E-05 | 0.016 | 1.75E-03 | 0.012 | 0.019 | 5.69E-19 | Egger | **4.55E-18** |

Here we present the best result for each trait considering different $r^2$ thresholds and the use of different GWAS datasets (Metabochip or jointGWAS), with multiple testing controlled by FDR. All effect sizes refer to the effect of having *lower* LDL-C. IVW: inverse variance weighted method; Egger, MR-Egger method; pleiotropy $p$ , p-value from test of unbalanced horizontal pleiotropy by the MR-Egger method; qval, q-values. Q-values < 0.05 are in bold; suggestive associations ( $0.05 < q < 0.2$ ) are in italics.
Please refer to Supplementary Table 2 for full results.

We observed significant associations of depressive symptoms and MDD. Lower LDL-C is causally linked to increased depressive symptoms (beta = 0.028, 95% CI 0.015 to 0.040, p = 1.04E-05) and higher risk of MDD (OR = 1.15, 95% CI 1.07 to 1.23, p = 9.31E-05) in this analysis. Both p-values passed the FDR control threshold at 0.05. We also observed potential causal relationship between lower LDL-C and reduced risk of schizophrenia (OR = 0.960, 95% CI 0.935 – 0.986, p = 2.81E-03). Other significant associations included lower LDL-C with longer sleep duration, lower platelet count and risk of celiac disease. We also observed a link between lower LDL-C with top 1% parental survival. For anorexia nervosa, the association was suggestive with q-value < 0.1. Low LDL-C was associated with reduced risk of anorexia. For Parkinson’s disease and forearm bone density, we did not observe any significant associations, suggesting that for these traits any causal relationships with statins, if present, may be to a greater extent attributed to HMG-CoA reductase inhibition than to a general lowering effect of LDL-C.

For most outcomes the MR estimates were based on the IVW approach, except for celiac disease and top 1% parental survival where the estimates were derived from MR-Egger due to significant horizontal pleiotropy. Full results are presented in Supplementary Table 2. Of note, for each trait under study, the directions of effects (provided that the result was at least nominally significant) were consistent despite different r^2^ thresholds or dataset (joint or Metabochip only) used.

We noted a very recently published study^17^ which employed MR to study the effects of statins, PCSK9 inhibitors and LDL-C on AD and PD risks. The study initially reported that lower LDL is causally associated with reduced AD risk (genetic variants in APOE excluded), using summary statistics from GLGC and International Genomics of Alzheimer’s Project (IGAP). Upon further analysis with a larger APOE gene region excluded, the authors revealed no significant casual relationships, which concurred with another recently published work in ophthalmology but used AD as a negative control^36^. We did not pursue further analyses here as it appeared that the causal links between LDL-C and AD (as revealed from MR) may be contributed mainly by APOE region variants. However, this does not exclude the possibility that other specific mechanisms (which may be diluted when other variants are included as instruments), such as HMGCR inhibition, may exert an effect on AD.

## DISCUSSION

In this study we have employed a Mendelian randomization approach to study the casual effects of genetically lowered LDL-C due to HMG-CoA reductase inhibition (analogous to the action of statins) on a large variety of traits. The analyses revealed several potential repositioning opportunities as well as adverse effects of statins. We shall discuss below associations showing at least nominal significance (*p* < 0.05) in our analyses.

### Depression, statins and LDL-C levels

One important observation from this study is a potential causal link between genetically lower LDL-C from HMG-CoA reductase inhibition and increased depressive symptoms. Consistent with this observation, we also detected causal links of lower LDL-C levels with elevated MDD risk and depressive symptoms. Although there was ~10% overlap in the two samples, they were still largely different, and it is noteworthy that we observed significant *inverse* relationships between LDL-C and depression risk or symptom scores.

A number of studies have been performed to elucidate the relationships between cholesterol levels (including LDL-C), statins and depression, although the results were mixed. You et al.^10^ provided a detailed review of the evidence linking depression with low cholesterol levels and statin use. For example, low serum cholesterol has been reported to increased rates of suicide and increased depressive symptoms in a number of reports (*e.g.* ref.^37-41^). Similarly, statin use has been suggested to be associated with elevated rates of depression (e.g. ref.^42-45^). The underlying mechanism remains to be elucidated, but some studies suggested that reduction of cholesterol may result in disruption of serotonin receptor functions^10^. For instance, Shrivastava et al. showed that chronic cholesterol depletion resulted in impaired ligand binding and G-protein coupling to serotonin 1A receptors^46^. Another possible mechanism is through reduced synthesis of neurosteroids^47^, which are important for normal brain functioning such as synaptogenesis^48^.

On the other hand, it has been postulated that the reduction of inflammatory responses by statins may contribute to antidepressant effects and some studies also found reduced depression risks among statin users^49^. For example, a recent meta-analysis of 3 randomized controlled trials (RCTs) (total *N* = 165) revealed efficacy of statins in treating depression, although one should caution against the small sample size and number of studies^50^. An earlier meta-analysis showed no significant differences in mental well-being between patients on statins or placebo, however there was high heterogeneity between the studies and psychological well-being were mainly assessed by self-report questionnaires^51^. There were also concerns of publications bias.

In the present study, we employed genetic instruments to model LDL-C levels, which reflect a life-long exposure to reduced LDL-C and it may not fully mimic the case of statin users. Therefore, it remains to be elucidated whether the observed effects on depressive symptoms from MR analyses can be directly translated to clinical practice. However, our results suggest that clinicians could be alerted to the possibility that development or exacerbation of depressive symptoms may be causally linked to prescription of statins or LDL-C lowering. For patients with risk factors for depression (e.g. family or past history of depression) who intend to receive or are receiving statins for prolonged periods, a greater awareness on mood changes and other psychiatric symptoms may be warranted.

Our results do not invalidate the possibility that statins may exert antidepressant properties under certain clinical scenarios. For example, significant results in RCTs^50^ imply that a relatively short-term course of statin to ameliorate inflammatory responses may be beneficial for certain depressive patients. Further studies are required to clarify how the duration of statin treatment and patient characteristics (e.g. age, sex, presence of pre-existing depressive risk factors, cardiometabolic profiles, concomitant use of other medications like antidepressants etc.) may be associated with worsening or improvement of depressive symptoms.

### Other psychiatric disorders and statins and LDL-C

For anorexia nervosa (AN), we observed relatively consistent results from single-SNP analysis of rs17238484 and rs12916, as well as from joint SNP analysis, that reduction of LDL-C from HMG-CoA reductase inhibition may reduce AN risk, although the statistical significance was not too strong. Concordantly, we also found suggestive associations between LDL-C and reduced risk of anorexia. Hypercholesterolemia, including raised LDL-C, is a well-known phenomenon in AN patients, although this association may appear paradoxical as these patients are generally malnutritioned and consume less fat than healthy individuals^52^. The underlying mechanism remains unclear. Although hypercholesterolemia is generally considered a consequence of AN, our results suggest that raised LDL (especially related to increased HMG-CoA reductase activity) may also be a casual risk factor for the disorder. Raised cholesterol synthesis has been suggested as a possible reason for hypocholesteremia in these patients^53^. Nevertheless, we are not aware of any studies on the effects of statins in AN patients. Clinical observational studies and preferably RCTs are required before one can conclude the role of LDL-C and benefits of statins in these patients.

We also observed some evidence that statins may reduce schizophrenia risk from individual SNP analysis of rs17238484. In support of this finding, we detected potential causal effects of LDL-C on schizophrenia risks in the same direction. Statin has been proposed as a novel treatment for the disorder, presumably based on its potential to ameliorate neuroinflammation^54^, and was tested in a small number of clinical trials. For instance, Vincenzi et al. reported improvement in the Positive and Negative Syndrome Scale (PANSS) from baseline to 6 weeks with pravastatin although the effect failed to maintain at 12 weeks^55^. In another study, there was preliminary evidence that simvastatin improved PANSS scores, although the results were not statistically significant^56^.

We found a slightly unexpected association of statin action with increased sleep duration in single-SNP MR analyses of rs17238484; interestingly, casual associations of reduced LDL-C with longer sleep duration was also detected. A limited number of observational studies suggested both short and long sleep durations are associated with lipid abnormalities^57,58^, although the current analyses could not detect non-linear relationships. With regards to effects of statins on sleep, observational studies have found statins may be associated with sleep disturbances^59^. However a recent meta-analysis of 5 RCTs (N = 231) showed that statins do not have adverse effects on sleep duration or efficiency^60^. A more recent study reported significantly *lower* rate of sleep disturbance in patients assigned atorvastatin than those assigned placebo in a double-blind RCT setting^61^.

### Neurological disorders and statins

Alzheimer disease (AD) is a common neurodegenerative disease for which few effective treatments are available. We found some evidence (*p* = 6.58e-3, *q* = 0.256) for statins to reduce AD risk from single-SNP MR analysis. The relationship between cholesterol lowering and statins with AD has been quite extensively studied. There is evidence from pre-clinical and clinical observational studies that statins may be able to prevent AD or mitigate the course of the disease^14,62,63^. The results from RCTs were more mixed, and no conclusive benefits of statins can be ascertained yet^64,65^. As with the study with other neuropsychiatric disorders, AD itself is a heterogeneous condition, and different patient characteristics (e.g. severity of illness, age, sex, comorbid illnesses etc.) in different studies may lead to mixed results. For Parkinson’s disease (PD), there was only one matching SNP but the MR result was nominally significant (*p* = 0.025, *q* = 0.274). There is some evidence from animal^66^ and observational studies (e.g. meta-analysis in ref.^67-69^) that statins may reduce PD risk.

A very recent study by Benn et al.^17^ also studied the *HMGCR* variant rs17238484 as a proxy for statin action in a Danish cohort. No associations were found with AD or PD, however the number of incident AD and PD cases was relatively small (1001 and 460 respectively), which may have limited the power to detect significant associations. We have found preliminary evidence for beneficial effects of statins in this study, but further studies are required to confirm the findings.

It is also worth mentioning that there are concerns about cognitive impairment as a side-effect of statins^70^. Besides studying AD, we also included a panel of cognitive test profiles as outcomes but did not find any significant associations. Although there is a chance of false negatives, this study and the previous work by Benn et al.^17^ appeared to show no causal relationship between statins and cognitive impairment.

#### Longevity and statins

We also revealed a potential polygenic association of *HMGCR* variants with extreme longevity (top 1% survival) and a highly significant association of LDL-C polygenic score with this trait (lowest p = 8.43e-9). The association is probably at least partially attributed to reduction in cardiovascular events. It remains unclear whether and by how much statins or reduction in LDL-C will benefit survival. A microsimulation study predicted that statin therapy is associated with an increased life expectancy of 0.3 years^71^. Our point OR estimate for extreme longevity was also relatively small for joint SNP analysis in *HMGCR* (OR = 1.024, 95% CI 0.999-1.049) and for LDL-C (OR = 1.016, 95% CI 1.012 to 1.019), although as explained above the estimate might be biased downwards.

#### Other possible associations

We observed a nominally significant association of increased forearm bone mineral density (BMD) with *HMGCR* variant. Statins are postulated to stimulate bone formation and hence improve BMD^72,73^. A recent meta-analysis of RCTs reported a small but significant benefit on BMD but no effects on fracture risk^74^. As we only detected a positive association for forearm BMD but not at two other sites (femoral neck and lumbar spine), the findings should be viewed with caution. Another nominal association was a potential decrease in platelet count with statins; we also reported a significant association of lower LDL-C with reduced platelet count. We did not find evidence from the literature that statins reduce platelet levels, although there might be an effect on reducing platelet aggregation^75^ and mean platelet volume^76^, which in turn might confer cardiovascular benefits. Limited evidence suggests a relationship between lower platelet counts and reduced cardiovascular risks^77^. Another nominal association was with celiac disease; again we did not find support from previous studies of any relationship between statins and celiac disease, except one study which showed that atorvastatin had no effect on gluten-induced production of inflammatory cytokines in intestinal biopsies^78^.

#### Negative findings

While a number of findings are negative, given the very widespread use of statins, these negative results are probably also of important public health significance. We did not find evidence that statins improve renal outcomes from MR analysis. A Cochrane review showed consistently lower mortality and cardiovascular events in chronic kidney disease (CKD) patients not requiring dialysis, but the effect on kidney function was inconclusive^79^. A more recent meta-analysis however showed beneficial effects of high-intensity statin therapy on estimated glomerular filtration rate (eGFR), while moderate- or low-intensity treatment did not show benefits^80^. Our negative findings may be due to statins only having an effect in a subgroup of patients with high-intensity therapy (which may be beyond the lipid-lowering effect of the studied genetic instruments). Alternatively, such beneficial effects may be due to off-target mechanisms. Similarly, no associations with cancers were found. There is currently no strong evidence to support chemopreventive potential of statins against most cancers^81,82^. Our results suggest that statins did not reduce or increase cancer risks. The current analysis, however, is limited by a relative lack of GWAS summary statistics available for cancers and relatively moderate sample sizes for some datasets. Finally, there were no associations found with autoimmune diseases. Parallel to this finding, there is lack of high-quality observational studies or RCTs demonstrating clear benefits of statins on the treatment of autoimmune disorders, although pre-clinical studies suggested statins may be useful for certain autoimmune diseases^83^. On the other hand, our results do not support the claim that statins may be a risk factor for autoimmune disorders in previous reports^84^.

We provided a discussion regarding our significant results with unadjusted *p* < 0.05 above, and noted many associations were supported by previous pre-clinical or clinical studies. Nevertheless, some of the findings could represent false positives (we also provided estimates of FDR in this report), and care must be taken before application of any results to clinical settings. Due to various limitations of an MR approach, we believe that the results did not yet provide confirmatory evidence for the repositioning potential or adverse effects of statins in actual practice, and further large-scale studies especially RCTs are needed to confirm the findings.

### Strengths and Limitations

This is the first study to employ a “phenome-wide” MR approach to explore drug repositioning opportunities and adverse side effects of statins and of a commonly used medication. The MR approach is analogous to a “naturalistic” RCT which is much less susceptible to confounding and reverse causation compared to clinical observational studies. This phenome-wide analysis provides an unbiased and comprehensive assessment of the therapeutic potential and side-effects of statins.

This study has several limitations. Firstly, as discussed previously, we employed genetic instruments to model the risk factor, reflecting a life-long exposure to altered LDL-C levels. Statin users typically receive the drug for a shorter period of time, and whether the effects of statins will be similar to those observed in the MR analyses requires further investigation via clinical studies. In addition, we focused on *HMGCR* variants which modelled the “on-target” effects of statins on HMG-CoA reductase (and its downstream pathways^85^). Off-targets effects might be missed. In a similar vein, we focused on a common mechanism of action by all statins, but it is possible that individual statins may exert more specific effects that are currently unknown. Pharmacokinetic properties of different statins, including ability to cross the blood-brain barrier, will need to be considered before applying specific statins for drug repositioning. Another potential limitation is that current GWAS mainly focus on identifying susceptibility variants for the development of disease, but few investigated the genetic basis underlying disease progression. As a result, MR analyses with drug target genes may be able to detect drugs potentially useful for disease prevention, but their effects on altering disease progression is less certain^86^. Nonetheless, a drug can be useful both in prevention and altering the disease course or preventing relapses, as is the case for statins for coronary heart disease^87^. A recent commentary^86^ provided a discussion on this issue. Also, the power of the MR analysis depends on the sample size of the GWAS studies, for several traits in our analysis (e.g. neurocognitive profiles and several cancers) the sample sizes are relatively modest and false negative results are possible.

Another potential source of bias is the collider bias, which in this study involves GWAS datasets of the UK Biobank (UKB). This issue has been recently discussed in Munafo et al.^88^. Briefly, there is evidence of “healthy volunteer bias” among UKB participants and the sample was healthier than the general UK population^89^. If an exposure and an outcome both influence the probability of participation, spurious associations between the two may result, as exposure and outcome are both conditioned on ‘participation’, or the collider in this case. In our current study, only a few GWAS involved the UK BioBank. For the GWAS on depressive symptoms^90^, ~58% of the sample was recruited from the UK BioBank. Although we did not have data on the rates of hyperlipidemia in UKB, a recent study reported much lower rates of cardiovascular diseases and risk factors (e.g. high blood pressure, diabetes) among UKB subjects^89^. It is probably reasonable to expect low LDL and low depressive symptoms are both linked to higher chance of participation in UKB; collider bias will lead to a positive association between low LDL and reduced depressive symptoms. Since we now observed causal relationships in the *opposite* direction, the findings cannot be explained by collider bias. In addition, a concordant causal link (lower LDL-C causing increased depression) was detected when we considered the PGC-MDD^91^ sample alone. Sleep duration was also extracted from UKB data, but there is no strong justification to believe that this variable has substantial impact on participation, nevertheless we cannot exclude possibility of collider bias. Yet another trait, top 1% survival in parents, may be more subject to this bias. Longevity in parents leads to longevity in offspring, and this factor along with low LDL may both be linked to participation in UKB, which may create a positive bias in the association between the two variables.

In summary, we showcased a phenome-wide MR approach to uncover repositioning opportunities and side-effects for statins. A number of findings, such as potential adverse effects on depressive symptoms and repositioning potential for several neuropsychiatric disorders, are worthy of further investigation. On the other hand, negative findings might also be of value due to the very widespread use of the drug. We hope that further preclinical experiments, clinical studies and RCTs will help to follow up and verify our findings.

## Acknowledgements

This study is partially supported by the Lo-Kwee Seong Biomedical Research Fund and a Direct Grant from the Chinese University of Hong Kong. We would like to thank the Hong Kong Bioinformatics Center for computing support.

## Author contributions

HCS designed, conceived and led the study. HCS performed the main data analysis, with assistance from CKLC and KZ. HCS interpreted the results and wrote the manuscript.

